# Feasibility Analyses and Experimental Confirmation of Dove Prism Based Dual-fiberscope Rotary Joint

**DOI:** 10.1101/2022.09.25.509388

**Authors:** Yuehan Liu, Hyeon-Cheol Park, Haolin Zhang, Xingde Li

## Abstract

Two-photon fluorescence microscopy has enjoyed its wide adoption in neuroscience. Head-mounted miniaturized fiberscopes offered an exciting opportunity for enabling neural imaging in freely-behaving animals with high 3D resolution. Here we propose a dual-fiberscope rotary joint based on a Dove prism, for enabling simultaneous two-photon imaging of two brain regions with two fiberscopes in freely-walking/rotating mice. Analytic proof has confirmed the key properties of a Dove prism. Feasibility analyses and proof-of-concept experimental results have demonstrated the feasibility of such a rotary joint for allowing two fiberscopes to rotate simultaneously while maintaining an excellent single-mode fiber-to-fiber coupling for the excitation femtosecond laser. Fiberscopes with a dual-probe rotary joint offer an exciting opportunity to explore neural network dynamics of multiple interconnected brain regions in freely-walking rotating animals.

## 1. INTRODUCTION

Brain activities involve neurons generating fast-propagating signals to encode and relay information within dynamic neural networks. Neuroscientists aspire to obtain access to such networks with a high spatiotemporal resolution, which will shed light on the fundamental working mechanisms of the brain. Optical imaging, particularly two-photon fluorescence (TPF) microscopy, has played a significant role in this endeavor [1, 2], which has exhibited multiple advantages such as high imaging resolution, 3D imaging capability [3], deeper penetration depth [4], and the ability to simultaneously excite multiple fluorophores with a single light source [5]. With the development of genetically encoded fluorescent calcium indicators (such as e.g., GCaMP), bench-top two-photon (2P) microscopy has become one of the key platforms for neural network imaging [6–8]. The past decade has witnessed many impressive progresses, from head-restrained benchtop microscopy virtual navigation to the developments of large FOV microscopy for neuron population imaging [9–11].

Another technological trend in this field is the miniaturization of imaging devices to enable real-time imaging of freely behaving rodents [12]. Our group has developed the first, fully integrated, fiber-optic scanning 2P endomicroscope for fluorescence and second harmonic generation imaging [13, 14]. By introducing a customized double-clad fiber (DCF) of a pure silica single-mode core, label-free *in vivo* 2P imaging at subcellular resolution has been achieved [15]. Along with a newly developed single-probe optoelectrical commutator (OEC), *in vivo* functional neural dynamics imaging in freely-behaving mice has been demonstrated [16–18]. In spite of all these exciting advances, tools for simultaneous 2P imaging over multiple brain regions in freely-behaving animals (e.g., rodents) are still lacking.

The capability of simultaneous imaging over multiple interconnected brain regions, along with the option for multicolor imaging, would provide an exciting opportunity for studying synergized functional connectivity of involved neural networks, and provide unprecedented insight into the neural circuit dynamics associated with various behaviors at population level with subcellular resolution. In addition, the freely-moving style would minimize the differences between experimentally controlled actions and natural behaviors, therefore, allowing precise examination of neural network functions. However, there are several bottlenecks: The large footprint (i.e., 15×9×21 mm^3^) of the state-of-the-art 2P miniscopes [12] makes it very challenging (if not impossible) to mount two 2P miniscopes over a very limited cortical area on a mouse head. In addition, the weight of two miniscopes would impose a prohibitively heavy burden (> 4.90 g) onto the mouse. One solution is to adopt the recently developed 2P fiberscopes which are extremely light (< 1 g for each fiberscope head) and compact (2-2.4 mm in diameter). However, an advanced fiber-optic rotary joint is needed for dual-probe imaging of freely walking/rotating rodents. Although a rotary joint for single-mode core-to-core optical coupling from one stationary source fiber to another one rotating probe fiber has been well-established, only very few vendors offer dual-fiber rotary joints at communication wavelengths (e.g., 1300-1500 nm) but with very poor coupling efficiency and rotational coupling variation (See Supplement 1). Due to a much smaller single-mode fiber core (~5 μm) at the 800-900 nm wavelength range, the coupling is even more challenging. To the best of our knowledge, currently such a dual-fiberscope rotary joint does not exist.

Here, we propose a dual-fiberscope rotary joint based on a Dove prism, allowing two fiberscopes to rotate simultaneously while maintaining an excellent single-mode fiber-to-fiber coupling for the excitation femtosecond laser from the stationary source fibers to the two fiberscopes. In this paper, we first present analytic proofs to confirm the key properties and working principle of a Dove prism based dual-probe rotary joint. We then report the feasibility of the dual-probe rotary joint using Ray-tracing analyses with the fabrication tolerance/error of key parameters of a Dove prism taken into account. Finally we demonstrate the initial proof-of-concept experimental results, confirming the feasibility of our proposed dual-probe rotary joint.

## 2. DESIGN AND METHODS

A critical component for the rotary joint to accommodate two fiberscopes is the Dove prism, which is a truncated right-angle prism with a base angle *α* (usually 45°). For a given material, the length and aperture size of the prism are designed based on the following Dove prism formula [19]:

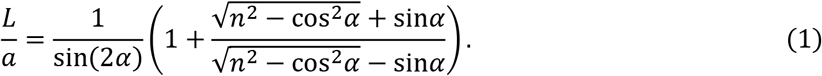

Here *L* is the length of the base (longest bottom face) of the prism, *a* is the side length of the prism cross section (or aperture), *α* is the base angle and *n* is the refractive index of the prism. Since the refractive *n* is wavelength dependent, the length *L* of a Dove prism needs to be specifically chosen for a given wavelength in order to achieve target performance. It is noted that a Dove prism can be used as an image inverter and rotator. We first define the rotation axis (RA) of a Dove prism as the axis parallel to the prism base and going through the center of the aperture (see Figure 1). A dove prism has several unique and very attractive optical properties. 1) For an incident beam parallel to the RA, the exit beam from the prism (after undergoing total internal reflection at the prism base) remains parallel to the RA; 2) For an incident beam parallel to the RA, the distance of the exit beam to the RA remains the same as the distance of the incident beam to the RA. This is crucial for fiber coupling of light since any lateral shift of the beam away from the RA will affect the coupling efficiency of the beam into a single-mode fiber (or the single-mode core of a DCF) [20]; 3) When the Dove prism is rotated by an angle *θ*, the exit beam (i.e., the image of the stationary incident beam parallel to the RA) rotates by an angle 2 *θ*. This means the rotation of a fiberscope can be compensated by rotating the Dove prism by a half angle so that the stationary incident beam can still be coupled into the rotated fiberscope after going through the half-angle rotated Dove prism; 4) The optical pathlength of any incident beams that are parallel with each other is a constant, which means a Dove prism does not introduce an optical pathlength difference among these beams. This is very favorable for 2P imaging since the material dispersion of the Dove prism for the incident femtosecond (fs) pulses can be conveniently pre-compensated, e.g., by using a grating-prism (GRISM) pair (for simultaneous compensation of the group velocity dispersion but for the third order dispersion as well) [16]. These key properties make a Dove prism a viable choice for a dual-fiberscope rotary joint. Analytic proofs according to geometric optics of these properties are provided in Supplement 2. Ray-tracing simulations by ZEMAX also confirm these properties.

**Figure 1.**
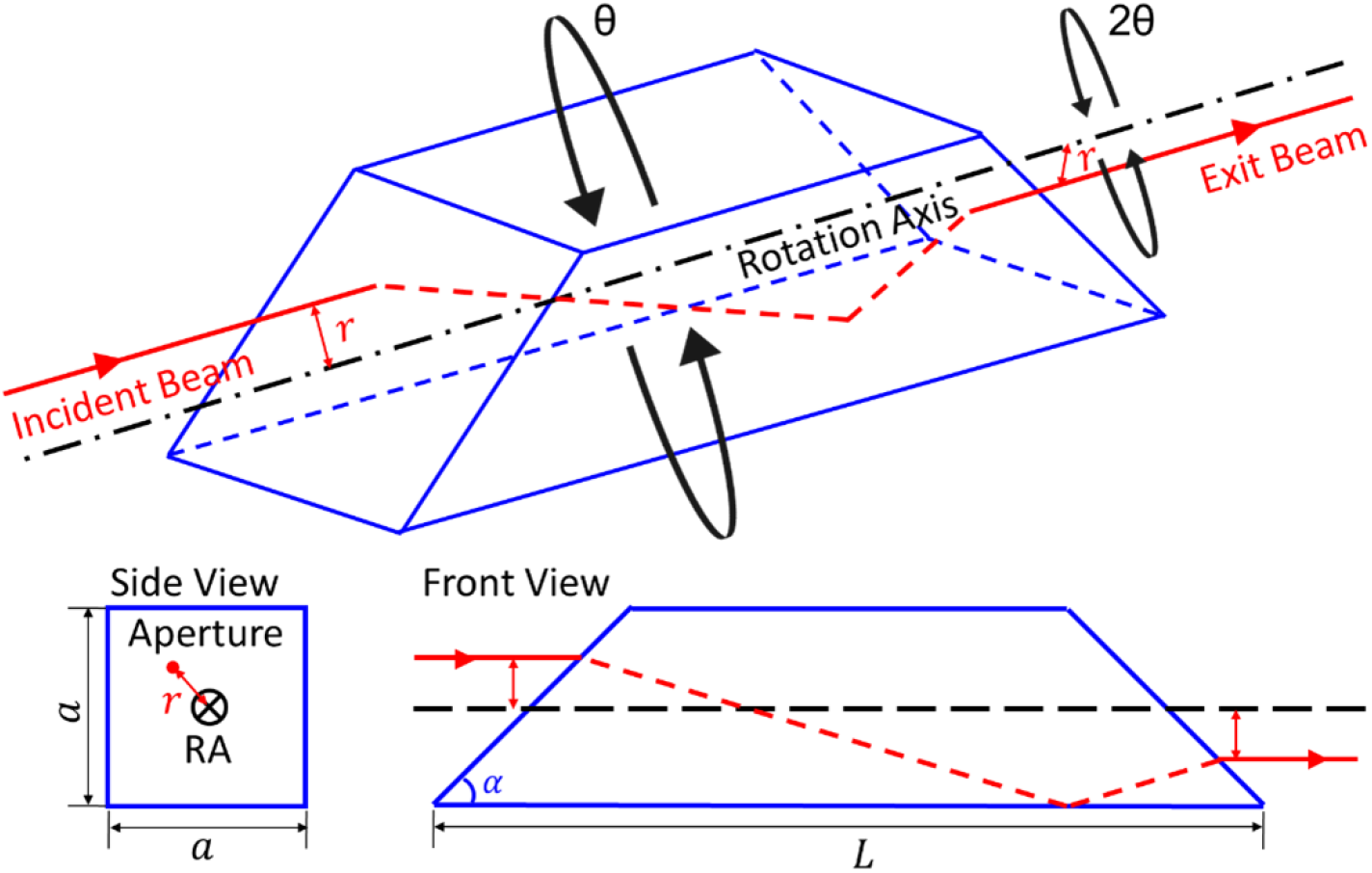
The Dove prism rotation axis (RA) is perpendicular to the prism aperture and also goes through the center of the aperture as shown in the side view as well as the front view.

The design schematic of a Dove prism based dual-fiberscope rotary joint is shown in Figure 2. In essence, two pairs of fiber-optic collimators (FCs) are used to couple light from the stationary source fibers to the rotating probe fibers (or fiberscopes). A Dove prism is sandwiched between FC1&2 and FC3&4. Note that the exit beam from the Dove prism rotates at twice the rate of the prism rotation. Therefore, the Dove prism and FC3&4 are mounted on two separate coaxial rotation shafts. Once the two pairs of FCs (FC1→FC4, FC2 →FC3) and the two rotation axes are precisely aligned, a fiberscope rotation angle 2*θ* can be compensated by *θ* rotation of the Dove prism, and the two incident beams can thus be efficiently coupled into the two fiberscopes through, the single-mode cores of the two DCFs in the fiberscopes [13]. The 2P fluorescence photons collected by the any of two fiberscopes (mainly through the large outer cladding of the DCF) can be separated by a dichroic mirror (DM) and then focused onto a photomultiplier tube (PMT) for detection (Figure 2). Although the fluorescence wavelength is different from the designed working wavelength (i.e., the 2P excitation wavelength) for the Dove prism and thus the TPF signals deviate from the excitation beam paths, the large detection area of PMT is good enough for detection. Here we consider GCaMP-based neural imaging as an example. According to Zemax simulations, the lateral shift of GCaMP fluorescence (around 525 nm) beam from the 920 nm excitation beam is less than 0.2 mm, which is much smaller than the photo cathode size in a PMT commonly used for 2P imaging.

**Figure 2.**
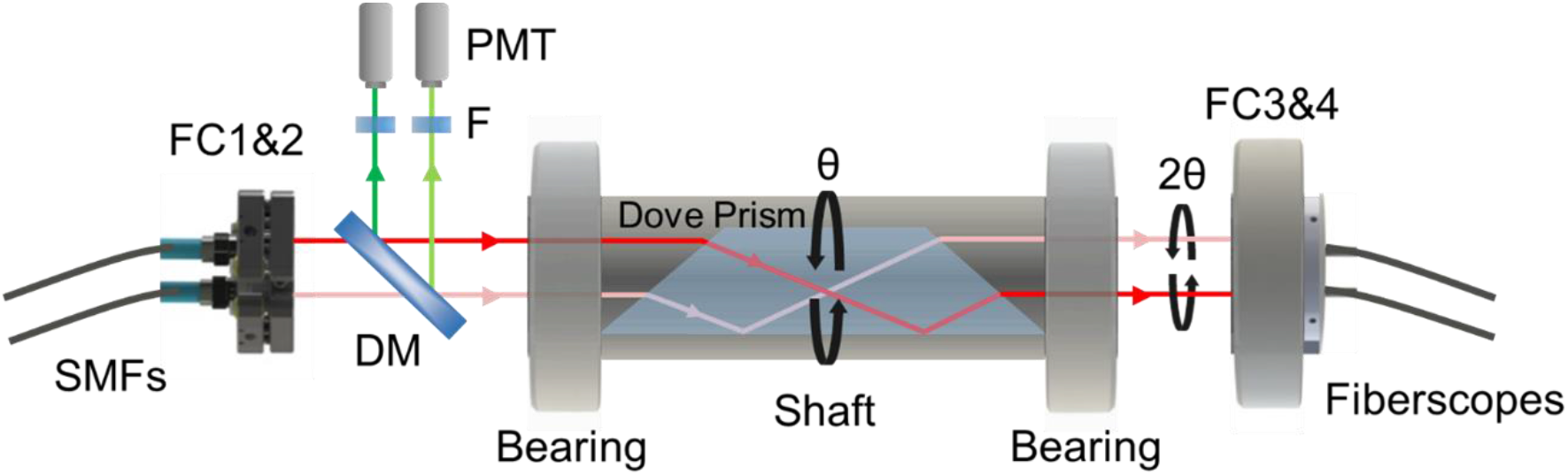
Schematic of a dual-fiberscope rotary joint based on a Dove prism. SMF: single-mode fiber (from the light source); FC: fiber collimator; DM: dichroic mirror; F: filter. The Dove prism is inserted into a hollow shaft which is mounted through two bearings. Two fs laser incident beams from the stationary FC1&2 go through the Dove prism and are coupled into rotary FC3&4 which are connected with two fiberscopes.

## 3. PRELIMINARY STUDIES AND RESULTS

Ideally, the rotary joint with a Dove prism should provide excellent stability in optical coupling efficiency at any rotational angle. However, misalignment of any optical components would result in optical throughput variation over rotation [20]. In addition, an imperfect Dove prism itself with manufacturing error/tolerance in geometry parameters (such as the length *L* and/or the base angle *α*) would also impact the coupling efficiency [21]. Furthermore, mismatch between the incident laser wavelength and the designed wavelength for the Dove prism will lead to small but non-negligible lateral beam shift, which will reduce the coupling efficiency and stability as well.

To investigate the feasibility of the dual-fiberscope rotary joint based on a Dove prism, proof-of-concept experiments have been conducted. The most critical parameter to test is the coupling efficiency stability for light coming from a stationary fiber collimator (i.e., FCsta in Figure 3a for the light from the laser) to the rotating fiber (i.e., the fiberscope) and a fiber coupler after going through a half-angle rotating Dove prism (see Figure 3a). Before a customized Dove prism with a proper length and aperture for a specific wavelength becomes available, we selected an off-the-shelf one (PS992, Thorlabs) which was intended for 675 nm light. We then chose a laser as the input light source available to us with a wavelength (668 nm) that is close to the Dove prism working wavelength (675 nm). As shown in Figure 3, on the stationary side, FCsta (CFC11P-B, Thorlabs) is connected to an x-y linear translation stage with a kinematic mount. On the rotary side, FCrot (F240APC-B, Thorlabs) is mounted on a rotary kinematic mount, whose optical axis can be precisely aligned parallel to its rotational axis. Once the two FCs and the Dove prism are well aligned, the incident beam (668 nm) coming out of a stationary single-mode fiber SMF1 could be effectively coupled into the single-mode fiber SMF2 at the rotary end with a minimum power fluctuation over rotation.

**Figure 3.**
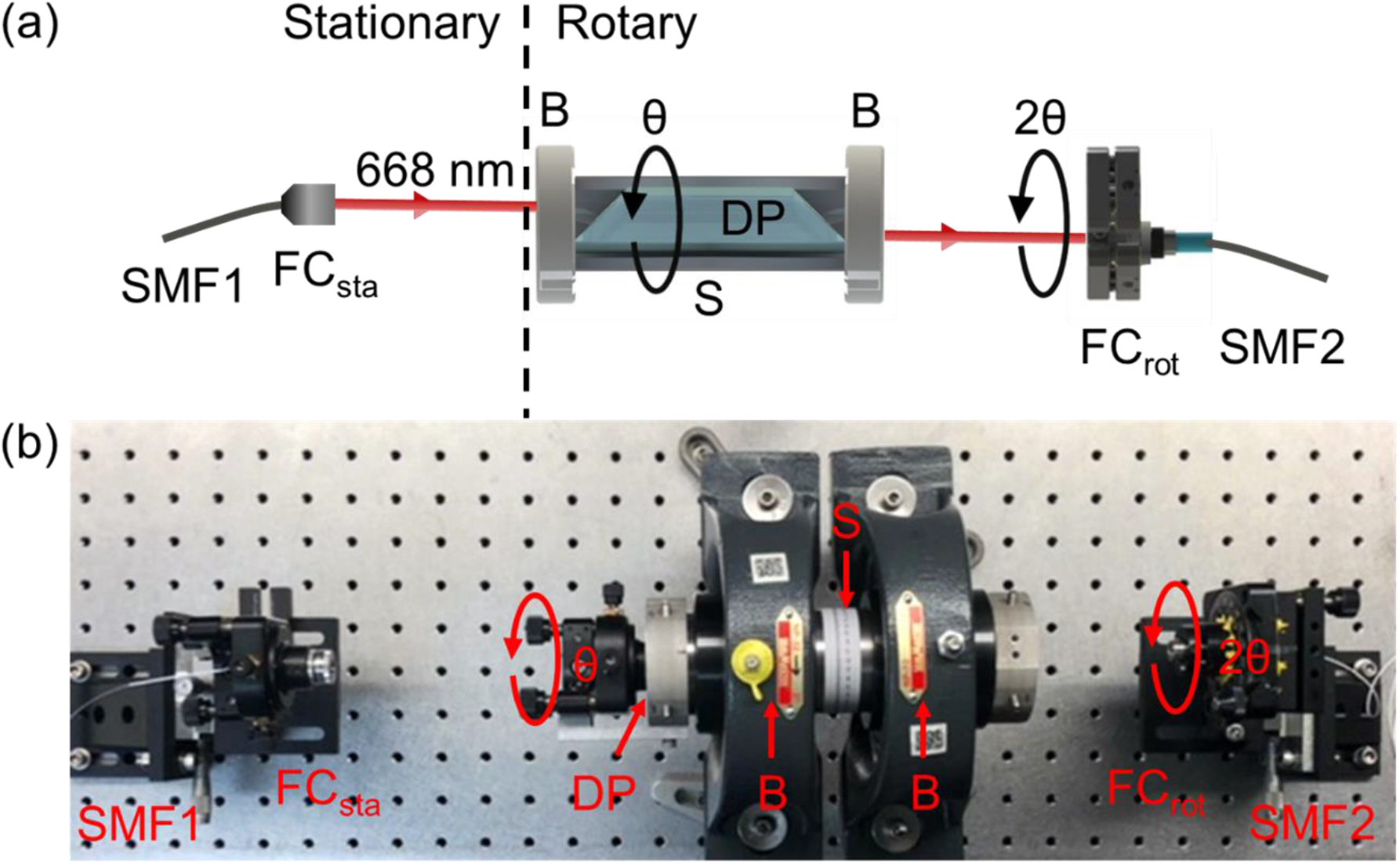
(a) Schematic and (b) Photograph of preliminary experimental setup for a Dove prism based rotary joint with one pair of fiber collimators (FC_sta_ & FC_rot_). DP: Dove prism; B: bearing; S: shaft.

We first performed quantitative Ray-tracing analyses using ZEMAX where the fabrication tolerance/error and wavelength mismatch for the off-the-shelf Dove prism were taken into account. The optical coupling efficiency of the FC pair (FCsta & FCrot) at 668 nm was calculated as a function of angular and lateral misalignment between two FCs. We concluded that to keep the coupling fluctuation below ±3% (which would not impact the analyses of dynamic neural activities for two-photon fiberscopy brain imaging of rodents [16]), the angular and lateral alignment tolerance over 360° rotation should be kept below ±2.4 mdeg and ±57.1 μm, respectively. Here, the Dove prism employed in our proof-of-concept experimentation has a fabrication tolerance/error (i.e., ±0.15 mm in length and ±0.05°in base angle). Taking into consideration both the wavelength mismatch (which is translated to a prism length mismatch) and the fabrication tolerance of the Dove prism, a maximum lateral shift for the exit beam reaches 79.0 um, corresponding to a normalized SMF coupling efficiency drop from an ideal 100.00% to 94.23% (see Figure 4a and b). This means the off-the-shelf Dove prism itself would cause approximately ±3% throughput fluctuation even.

**Figure 4.**
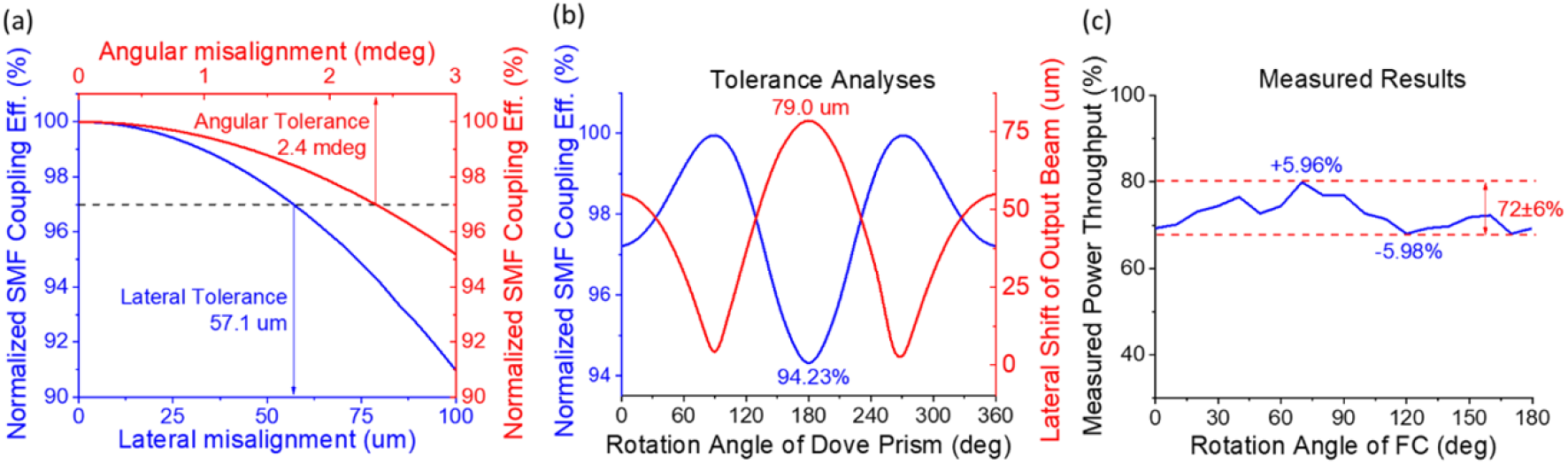
Preliminary feasibility studies of the performance of a Dove prism based rotary joint. (a) Normalized SMF coupling efficiency versus lateral and angular misalignment based on quantitative Raytracing analyses using Zemax. (b) Fabrication tolerance analyses by quantitative Ray-tracing. Blue curve: normalized SMF coupling efficiency with a Dove prism sandwiched between two FCs. Red curve: lateral shift of output beam caused by wavelength mismatch (for a Dove prism PS992 designed for 675 nm with a laser used in experiments at 668nm) and fabrication errors of the Dove prism. (c) The measured normalized optical throughput fluctuation over 180° rotation of FCrot.

Encouraged by the Ray-tracing analyses, we proceeded with experimental testing. The laser power throughput from SMF1 to SMF2 was measured to be 72% which is excellent. A better than ±6% relative fluctuation was achieved in the throughput (or coupling efficiency) over 180° rotation of FCrot (accompanied by the compensating half angle rotation of the Dove prism over 90°) (see Figure 4). It is noted that only 180° (rather than 360°) rotation of the FC_rot_ is needed for testing owing to the rotational symmetry. This excellent coupling or throughput stability was obtained even with non-precision bearings and a non-tight-fit housing shaft available in our lab. It is noticed that the rotational fluctuation in the coupling efficiency is about 2X as large as the simulation results, and this larger fluctuation was due to the imperfect off-the-lab-shelf mechanical components (two ball bearings and rotating shaft) as well as the mismatch between the intended wavelength for the generic Dove prism and the laser wavelength available to us. The measured throughput fluctuation was a lightly less than ±6% (see Figure 4c). This translates to ±12% rotation-induced fluctuation in the fluorescence signal (ΔF/F) during two-photon imaging of neural dynamics, which is still considered acceptable since the relative dynamic change of neural activity related two-photon fluorescence signal ΔF/F is generally greater than 50%.

## 4. CONCLUSION AND DISCUSSIONS

We have quantitatively (using Ray-tracing) and experimentally demonstrated the feasibility of a Dove prism based rotary joint. We have analytically proved the unique optical properties of a Dove prism suited for a dual-probe rotary joint. Quantitative Ray-tracing analyses support the feasibility of such a dual-probe rotary joint. We further performed proof-of-concept experiments. Even without the use of precision mechanical components and a prism of an unmatched length for the test laser wavelength, we were still able to achieve a high coupling efficiency (~72%) and a fairly small rotational fluctuation (±6%).

Considering the quadratic dependence of the 2P fluorescence on the excitation intensity 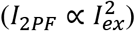, a variation in excitation laser intensity would induce a higher variation in the fluorescence signal. Assuming the target rotation-induced fluorescence fluctuation is less than 10%, the acceptable excitation throughput variation shall be maintained less than 5% over rotation. If needed, a smaller rotational variation in coupling efficiency can be achieved by slightly sacrificing the coupling throughput, which can be compensated by slightly increasing the input power from the laser.

For two-photon imaging of GCaMP based neural activities, a longer fs excitation wavelength at 920nm will be used. The corresponding single-mode fiber core of the fiberscopes will be about 40% larger than the SMF for 668 nm light used in the above experiments. The increased core diameter will help achieve stable coupling. The use of a customized Dove prism designed for the exact wavelength of the excitation light (i.e., 920nm) would reduce the rotational fluctuation in the coupling efficiency. In addition, the coupling stability can also be improved by using precision bearings (as opposed to the ones we have in the lab) and a shaft with a proper diameter for tight fit to the bearings’ inner diameter.

Although in the above experimentation we only considered one fiberscope (connected with FC_rot_) in the above experiments, the same exercise can expand to a dual-probe configuration. Smaller kinematic mounts can be used for the FCs or beam steering mirrors can be used in the beam paths to avoid potential beam blocking by the mechanical components when two fiberscopes are connected to the dual-probe rotary joint. Such a Dove-prism based rotary joint would enable for the first time simultaneous 2P imaging of two brain regions in freely-walking/rotating mice. In principle, our design is not only restricted to neuroimaging of rodents. Owing to the twist-free operation, it can be applied to non-human primates like rhesus macaque and marmoset. We believe the system will open a new avenue for exploring neural network dynamics of multiple interconnected brain regions associated with various behaviors.

## Supporting information

Supplement 1: Table Survey of off-the-shelf dual-channel fiber-optic rotary joints

Supplement 2: Analytic Proof of Dove Prism Properties with Geometric Optic

## ACKNOWLEDGEMENT

The authors are grateful for the partial support of this work by the Bisciotti Foundation (Li and Park) and the National Institutes of Health under a grant R35CA209960 (Bhujwalla).

## SUPPLEMENTAL MATERIALS

**Supplement 1:** Table Survey of off-the-shelf dual-channel fiber-optic rotary joints

**Supplement 2:** Analytic Proof of Dove Prism Properties with Geometric Optic

## REFERENCES

[1] W. Denk, J. H. Strickler, and W. W. Webb, “Two-photon laser scanning fluorescence microscopy,” Science 248, 73–76 (1990).

[2] K. Svoboda, and R. Yasuda, “Principles of two-photon excitation microscopy and its applications to neuroscience,” Neuron 50, 823–839 (2006).

[3] W. R. Zipfel, R. M. Williams, and W. W. Webb, “Nonlinear magic: multiphoton microscopy in the biosciences,” Nature biotechnology 21, 1369–1377 (2003).

[4] F. Helmchen, and W. Denk, “Deep tissue two-photon microscopy,” Nature methods 2, 932–940 (2005).

[5] C. Xu, W. Zipfel, J. B. Shear, R. M. Williams, and W. W. Webb, “Multiphoton fluorescence excitation: new spectral windows for biological nonlinear microscopy,” Proceedings of the National Academy of Sciences 93, 10763–10768 (1996).

[6] L. Tian, S. A. Hires, T. Mao, D. Huber, M. E. Chiappe, S. H. Chalasani, L. Petreanu, J. Akerboom, S. A. McKinney, and E. R. Schreiter, “Imaging neural activity in worms, flies and mice with improved GCaMP calcium indicators,” Nature methods 6, 875–881 (2009).

[7] R. P. Barretto, T. H. Ko, J. C. Jung, T. J. Wang, G. Capps, A. C. Waters, Y. Ziv, A. Attardo, L. Recht, and M. J. Schnitzer, “Time-lapse imaging of disease progression in deep brain areas using fluorescence microendoscopy,” Nature medicine 17, 223–228 (2011).

[8] W. Yang, and R. Yuste, “In vivo imaging of neural activity,” Nature methods 14, 349–359 (2017).

[9] D. A. Dombeck, A. N. Khabbaz, F. Collman, T. L. Adelman, and D. W. Tank, “Imaging large-scale neural activity with cellular resolution in awake, mobile mice,” Neuron 56, 43–57 (2007).

[10] D. A. Dombeck, C. D. Harvey, L. Tian, L. L. Looger, and D. W. Tank, “Functional imaging of hippocampal place cells at cellular resolution during virtual navigation,” Nature neuroscience 13, 1433–1440 (2010).

[11] J. N. Stirman, I. T. Smith, M. W. Kudenov, and S. L. Smith, “Wide field-of-view, multi-region, two-photon imaging of neuronal activity in the mammalian brain,” Nature biotechnology 34, 857–862 (2016).

[12] W. Zong, R. Wu, S. Chen, J. Wu, H. Wang, Z. Zhao, G. Chen, R. Tu, D. Wu, and Y. Hu, “Miniature two-photon microscopy for enlarged field-of-view, multi-plane and long-term brain imaging,” Nature methods 18, 46–49 (2021).

[13] M. T. Myaing, D. J. MacDonald, and X. Li, “Fiber-optic scanning two-photon fluorescence endoscope,” Optics letters 31, 1076–1078 (2006).

[14] Y. Zhang, M. L. Akins, K. Murari, J. Xi, M.-J. Li, K. Luby-Phelps, M. Mahendroo, and X. Li, “A compact fiberoptic SHG scanning endomicroscope and its application to visualize cervical remodeling during pregnancy,” Proceedings of the National Academy of Sciences 109, 12878–12883 (2012).

[15] W. Liang, G. Hall, B. Messerschmidt, M.-J. Li, and X. Li, “Nonlinear optical endomicroscopy for label-free functional histology in vivo,” Light: Science & Applications 6, e17082–e17082 (2017).

[16] A. Li, H. Guan, H.-C. Park, Y. Yue, D. Chen, W. Liang, M.-J. Li, H. Lu, and X. Li, “Twist-free ultralight two-photon fiberscope enabling neuroimaging on freely rotating/walking mice,” Optica 8, 870–879 (2021).

[17] H. Guan, D. Li, H.-c. Park, A. Li, Y. Yue, Y. A. Gau, M.-J. Li, D. E. Bergles, H. Lu, and X. Li, “Deep-learning two-photon fiberscopy for video-rate brain imaging in freely-behaving mice,” Nature Communications 13, 1534 (2022).

[18] H. Guan, W. Liang, A. Li, Y.-T. A. Gau, D. Chen, M.-J. Li, D. E. Bergles, and X. Li, “Multicolor fiber-optic two-photon endomicroscopy for brain imaging,” Optics Letters 46, 1093–1096 (2021).

[19] H. Sar-El, “Revised Dove prism formulas,” Applied optics 30, 375–376 (1991).

[20] C.-H. Chou, R.-J. Chen, H.-Y. Tsai, K.-C. Huang, and C.-C. Yang, “Optimization of Coupling Efficiency of Fiber Optic Rotary Joint by Ray Tracing,” in PHOTOPTICS (2020), pp. 71–75.

[21] D. Jia, C. Yu, W. Jing, H. Zhang, and Y. Zhang, “Effect of misalignment on rotating coupling efficiency of the Dove prism,” in Advanced Materials and Devices for Sensing and Imaging III(SPIE2008), pp. 128–136.

