## Supplement 1: Table Survey of off-the-shelf dual-channel fiber-optic rotary joints for "Feasibility Analyses and Experimental Confirmation of Dove Prism Based Dual-fiberscope Rotary Joint"

**Table**: Survey of off-the-shelf dual-channel fiber-optic rotary joints

| Vendor | Model | Loss (dB) | Rotary Variation (dB) | Rotary Variation (%) | Wavelength (nm) |
|---|---|---|---|---|---|
| FOCAL | 292 | 4.5 | 1 | 21 | 850/1310/1550 |
| Princetel | MJ2-131-62 | 4 | $\pm 1$ | $\pm 21$ | 850/1310/1550 |
| SENRING | FO209 | 4 | 2 | 37 | 1310/1550 |
| JINPAT | LPFO-02A | 5 | 2 | 37 | 850/1550 |
