## Supplement 2: Analytic Proof of Dove Prism Properties with Geometric Optic for "Feasibility Analyses and Experimental Confirmation of Dove Prism Based Dual-fiberscope Rotary Joint"

### Analytic Proof of Dove Prism Properties with Geometric Optic

Dove prism is a reflective prism which is used to invert an image or rotate an image in a rotating optical system [1]. It has several unique optical properties, allowing for the use of a Dove prism (as a critical element) in a dual-fiberscope rotary joint, which enables two fiberscopes to rotate simultaneously while maintaining an excellent single-mode fiber-to-fiber coupling. The key optical properties of a Dove prism are summarized below:

1) For an incident beam parallel to the RA, the exit beam from the prism (after undergoing total internal reflection at the prism base) remains parallel to the RA; 2) For an incident beam parallel to the RA, the distance of the exit beam to the RA remains the same as the distance of the incident beam to the RA. This is crucial for fiber coupling of light since any lateral shift of the beam away from the RA will affect the coupling efficiency of the beam into a single-mode fiber (or the single-mode core of a DCF) [2]; 3) When the Dove prism is rotated by an angle $\theta$, the exit beam (i.e., the image of the stationary incident beam parallel to the RA) rotates by an angle $2\theta$. This means the rotation of a fiberscope can be compensated by rotating the Dove prism by a half angle so that the stationary incident beam can still be coupled into the rotated fiberscope after going through the half-angle rotated Dove prism; 4) The optical pathlength of any incident beams that are parallel with each other is a constant, which means a Dove prism does not introduce an optical pathlength difference among these beams. This is very favorable for 2P imaging since the material dispersion of the Dove prism for the incident femtosecond (fs) pulses can be conveniently pre-compensated, e.g., by using a grating-prism (GRISM) pair (for simultaneous compensation of the group velocity dispersion but for the third order dispersion as well) [3].

Property 1 is obvious according to ray optics [4]. Here, we provide analytic proof for the other properties. We start with a Dove prism with a commonly used base angle $\alpha = 45°$, and then expand the conclusion to an arbitrary base angle. The schematic of a Dove prism of a square cross section (also called aperture) $A = a \times a$ and a base length $L$ is shown in Figure S1. Here, four planes are defined: the vertial input plane, 45-deg entrance surface, 45-deg exit surface, and the vertical output plane. The intersection points of the optical beam with these four planes are $(x_1, y_1, z_1)$, $(x_2, y_2, z_2)$, $(x_3, y_3, z_3)$, and $(x_4, y_4, z_4)$, respectively. (Define the rotation axis of a Dove prim as in the text). The rotation angle of the incident beam about the rotation axis is denoted as $\theta$ (assuming it is initially on the positive x-axis and rotation is along the clockwise direction).

For a Dove prism to work properly in many applications including the current intended use in a dual-probe rotatory joint, the base length $L$ and the aperture size $a$ of a Dove prism needs to satify the following equation (1):

$$\frac{L}{a} = \frac{1}{\sin(2\alpha)}\left(1 + \frac{\sqrt{n^2 - \cos^2\alpha} + \sin\alpha}{\sqrt{n^2 - \cos^2\alpha} - \sin\alpha}\right) \xrightarrow{\alpha=45°} 1 + \frac{\sqrt{n^2 - \frac{1}{2}} + \frac{\sqrt{2}}{2}}{\sqrt{n^2 - \frac{1}{2}} - \frac{\sqrt{2}}{2}}, \tag{1}$$

where *n* is the wavelength-deendent refratctive index of the Dove prism.

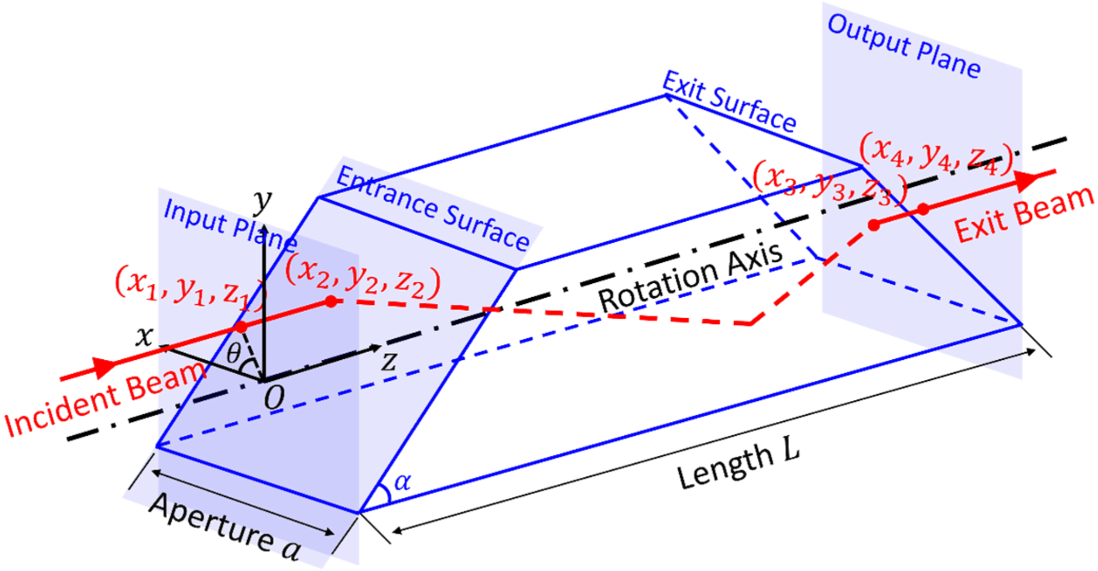


**Figure S1.** Schematic and parameters of a Dove prism. Reference planes and the intersection points of the incident beam with those reference planes are also explicitly labeled on the schematic.

Without losing generality, we consider $y_1 > 0$ (i.e., the incident beam is above the rotation axis).

**Proof of Property 2:**

The distance between the incident beam and the rotation axis is denoted as $r$. At the input plane, the beam is at $(x_1, y_1, z_1) = (r\cos\theta, r\sin\theta, 0)$. Since the beam is parallel to the z-axis, the incident point of the light on the entrance surface of the Dove prism can be found $(x_2, y_2, z_2) = \left(r\cos\theta, r\sin\theta, \frac{a}{2} + r\sin\theta\right)$.

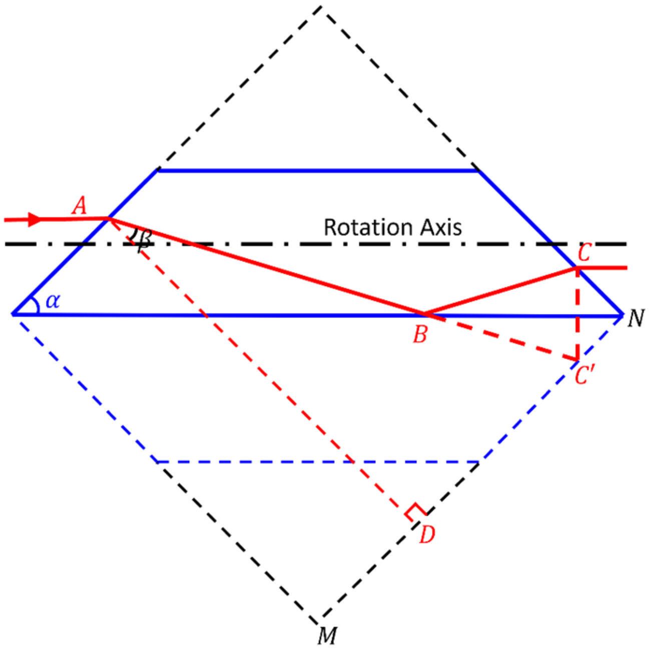


**Figure S2.** Optical path of a light beam parallel with rotation axis in a Dove prism ($\alpha = 45°$).

The coordinates of the beam at the exit surface of the prism can be found according to geometric optics.

On Figure S2 and according to Snell's Law, we have

$$n_0\sin\left(\frac{\pi}{2}-\alpha\right)=n_D\sin\beta, \tag{2}$$

where $n_0$ is the refractive index of air, $n_D$ is the refractive index of Dove prism, and $\beta$ is the refractive angle of the beam after the entrance surface.

To facilitate visualization, we introduce some auxiliary lines (indicated with dashed lines in Figure S2) where the prism base becomes the diagonal of the dashed square. Obviously, the graphic is symmetrical along the base of the Dove prism. From the geometric relationship, we have:

$$MD=\frac{\sqrt{2}}{2}a+\sqrt{2}y_1=\frac{\sqrt{2}}{2}a+\sqrt{2}r\sin\theta, \tag{3}$$

$$DC'=\frac{\sqrt{2}}{2}L\tan\beta, \tag{4}$$

$$C'N=\frac{\sqrt{2}}{2}L-\frac{\sqrt{2}}{2}a-\sqrt{2}r\sin\theta-\frac{\sqrt{2}}{2}L\tan\beta, \tag{5}$$

$$\frac{CC'}{2}=\frac{\sqrt{2}}{2}\left(\frac{\sqrt{2}}{2}L-\frac{\sqrt{2}}{2}a-\sqrt{2}r\sin\theta-\frac{\sqrt{2}}{2}L\tan\beta\right)=\frac{L}{2}(1-\tan\beta)-\frac{1}{2}a-r\sin\theta, \tag{6}$$

$$y_3=-\left(\frac{a}{2}-\frac{CC'}{2}\right)=\frac{L}{2}(1-\tan\beta)-a-r\sin\theta. \tag{7}$$

Therefore, the coordinates of the beam are $(x_3,y_3,z_3)=\left(r\cos\theta,\frac{L}{2}(1-\tan\beta)-a-r\sin\theta,\sqrt{2}L-\frac{a}{2}-y_3\right)$ on the exit surface of the Dove prism. According to Property 1, the exit beam is parallel to the rotation axis. Then we have $x_4=x_3$, $y_4=y_3$ and $(x_4,y_4,z_4)=\left(r\cos\theta,\frac{L}{2}(1-\tan\beta)-a-r\sin\theta,\sqrt{2}L\right)$ on the output plane. Note that the exit beam and the incident beam travel in the same plane (plane of incidence) so that $x_4=x_3=x_2=x_1$.

The distance of the exitbeam to the rotation axis (i.e., z axis) can be calculated and simplified:

$$\sqrt{x_4^2+y_4^2}=\left(r^2+2ar\sin\theta+a^2+\frac{L^2}{4}\cdot\frac{4a^2}{L^2}-La\cdot\frac{2a}{L}-2ar\sin\theta\right)^{\frac{1}{2}}=r, \tag{8}$$

where the relationship between $L$ and $a$ given by the Dove prism formula Eq. 1 and the parameter $\beta$ given in Eq. 2 have been used to simply the above expression. We find the exit beam, which is parallel to the ortation axis, remain the distance to the rotation axis as the incident beam during the relative rotation between the incident beam and the Dove prism.

**Proof of Property 3:**

Furthermore, since $x_4=x_1$, then $|y_4|=y_1$, $\theta'=\theta$.

This means when the Dove prism rotates around the rotation axis with a rotation angle of $\theta$, the output light beam rotates twice the angle ($\theta + \theta' = 2\theta$).

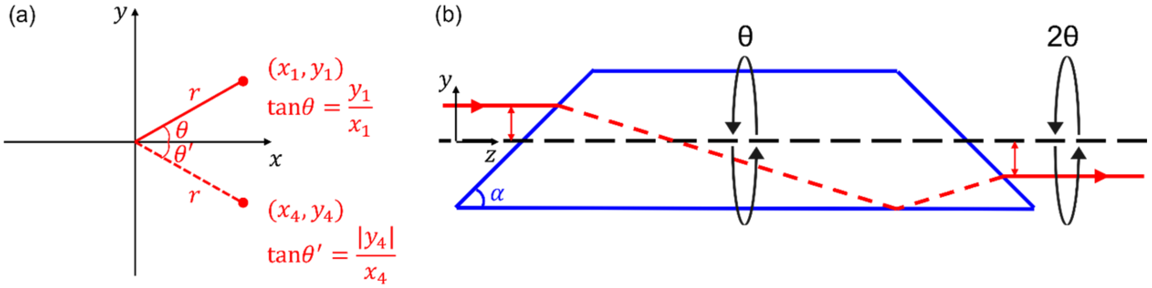


**Figure S3.** When the Dove prism rotates around the rotation axis with a rotation angle of $\theta$, the output light beam rotates twice the angle $2\theta$.

Furthermore, by substituting Eqs. (1) and (2) into Eq. (7), we find $y_4 = y_3 = -r\sin\theta = -y_1$. Since $x_4 = x_1 = rcos\theta$, we can easily find the exit beam rotates by the same angular amount but in the opposite direction. As shown inFigure S3, $\theta' = \theta$. This means when the Dove prism rotates around the rotation axis by an angle of $\theta$ relative to the incident beam, the exit beam rotates twice the angle ($\theta + \theta' = 2\theta$).

**Proof of Property 4:**

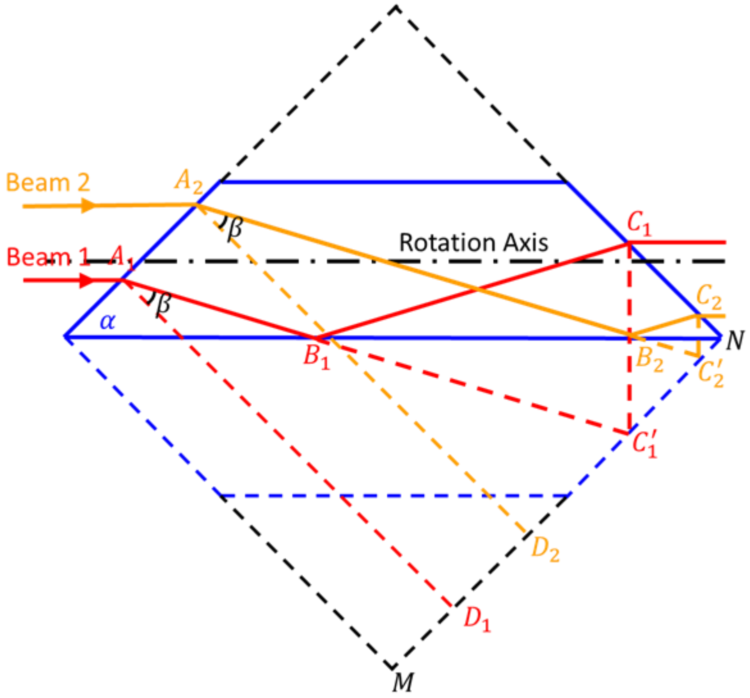


**Figure S4.** Optical pathlengths of beams parallel to each other are the same.

As shown in Figure S4, for two beams parallel to each other, we have:

Optical Pathlength for Beam 1: $A_1B_1C_1 = A_1B_1C_1' = \frac{A_1D_1}{\cos\beta} = \frac{\sqrt{2}L}{2\cos\beta}$, and

Optical Pathlength for Beam 2: $A_2B_2C_2 = A_2B_2C_2' = \frac{A_2D_2}{\cos\beta} = \frac{\sqrt{2}L}{2\cos\beta}$.

So the optical pathlengths of these beams within the prism are the same.

Please note that the above proofs are for a Dove prism of a 45° base angle ($\alpha = 45°$). The conclusions remain for an arbitrary base angle except that the expressions of the coordinates of $(x_4, y_4, z_4)$ are slightly more complicated.

### REFERENCES


[1] M. J. Padgett, and J. P. Lesso, "Dove prisms and polarized light," Journal of Modern Optics **46**, 175-179 (1999).

[2] C.-H. Chou, R.-J. Chen, H.-Y. Tsai, K.-C. Huang, and C.-C. Yang, "Optimization of Coupling Efficiency of Fiber Optic Rotary Joint by Ray Tracing," in *PHOTOPTICS*(2020), pp. 71-75.

[3] A. Li, H. Guan, H.-C. Park, Y. Yue, D. Chen, W. Liang, M.-J. Li, H. Lu, and X. Li, "Twist-free ultralight two-photon fiberscope enabling neuroimaging on freely rotating/walking mice," Optica **8**, 870-879 (2021).

[4] E. Hecht, *Optics* (Pearson Education India, 2012).
